# Simulated climate change causes asymmetric responses in insect life history timing potentially disrupting a classic ecological speciation system

**DOI:** 10.1101/2022.12.06.519222

**Authors:** Alycia C. R. Lackey, Pheobe M. Deneen, Gregory J. Ragland, Jeffrey L. Feder, Daniel A. Hahn, Thomas H. Q. Powell

**Author notes:** **Corresponding Author:** Thomas H. Q. Powell, Department of Biological Sciences, Binghamton University (State University of New York), Binghamton, NY 13902, USA. **Author Statement:** THQP, JLF, GJR, & DAH conceived of the study, THQP and ACRL designed the experiment, THQP, ACRL, and GJR made the field collections, ACRL, PMD, and THQP executed the experiment, ACRL and PMD analyzed the data, the manuscript was written by THQP and ACRL with input from all authors. **Data accessibility:** Data from the study (all information collected about individual puparia across the experiment) will be submitted to DRYAD upon publication.

## Abstract

Climate change may alter phenology within populations with cascading consequences for community interactions and on-going evolutionary processes. Here, we measured the response to climate change in two sympatric, recently diverged (~170 years) populations of *Rhagoletis pomonella* flies specialized on different host fruits (hawthorn and apple) and their parasitoid wasp communities. We tested whether warmer temperatures affect dormancy regulation and its consequences for synchrony across trophic levels and temporal isolation between divergent populations. Under warmer temperatures, both fly populations developed earlier. However, warming significantly increased the proportion of maladaptive pre-winter development in apple, but not hawthorn, flies. Parasitoid phenology was less affected, potentially generating ecological asynchrony. Observed shifts in fly phenology under warming may decrease temporal isolation, potentially limiting on-going divergence. Our findings of complex sensitivity of life-history timing to changing temperatures predict that coming decades may see multifaceted ecological and evolutionary changes in temporal specialist communities.

## INTRODUCTION

A major challenge in predicting the ecological and evolutionary consequences of climate change is understanding how responses within populations affect community interactions. Within populations, common responses to climate change include shifts in geographic distributions and phenology (Walther et al. 2002, Root et al. 2003, Parmesan 2006, Bradshaw & Holzapfel 2008, Cohen et al. 2018). Differential local adaptation, reduced gene flow, and associated genetic differentiation can result in population differences in life history and phenological responses (Forrest & Miller-Rushing 2010). When interacting species respond differently to climate change, phenological mismatches are likely (Renner & Zohner 2018, Rudolf 2019, Visser & Gienapp 2019), especially for specialists or when temporal windows for interaction are brief.

Life history timing is a crucial axis of ecological adaptation for temperate organisms. Different species rely on different sets of environmental cues (e.g., from photoperiod, temperature, aridity, and other organisms) to regulate life history timing to withstand unfavorable conditions and synchronize life cycles with ephemeral resources (Tauber & Tauber 1976, Williams et al. 2017, Wilsterman et al. 2021). Adaptive timing of life history, such as initiation of dormancy, involves the integration of a series of environmental cues (Denlinger 2002; Emerson et al. 2009). Thus, the ramifications of altered seasonality may be multifaceted, and phenological effects on populations may ripple through entire communities.

Studies of phenological change over recent decades of anthropogenic climate change have shown that altered seasonality of populations may already be driving ecological asynchrony (Parmesan 2006). However, robust experimental tests examining predicted future climatic conditions face tradeoffs between biological realism and experimental precision (Diamond 1986). While *in situ* warming experiments where temperature is manipulated in the field offer potentially powerful approaches (e.g., Hoffman et al. 2010; Diamond et al. 2016; Wagdymar et al. 2015)., inferences from these studies may be hampered by our ability to impose treatments on biologically relevant spatial and temporal scales across ecological communities. Thus, climate-induced shifts may be experimentally shown to alter phenology as well as evolutionary and ecological processes within populations. However, they may miss important consequences of these altered phenotypes on interactions between species across trophic levels in communities. Moreover, as divergent life history timing can promote temporal (allochronic) isolation between populations adapted to different seasonal regimes (Taylor and Friesen 2017), the evolutionary effects of climate change may facilitate or hinder speciation and thus the origin of biodiversity.

Here, we used an ideal system for testing the potential for ecological and evolutionary effects of climate change on temporal specialist insect populations that allowed us to impose realistic climate manipulations in the laboratory across two trophic levels. The Tephritid fruit fly *Rhagoletis pomonella* is a textbook example of ecological speciation-in-action (Berlocher & Feder 2002, Dres & Mallet 2002, Coyne & Orr 2004) and a model for rapid seasonal adaptation (Dowle et al. 2020; Powell et al. 2020). A recently derived, partially reproductively isolated host race of these flies evolved after shifting to domesticated apples (*Malus pumila*) from ancestral downy hawthorn fruit (*Crataegus mollis*) ~170 years ago, after apples were introduced to North America ~400 years ago (Bush 1969, Feder et al. 1988, Michel et al. 2010). Both populations are monophagous specialists, with a single generation per year and life cycles synchronized to the availability of ripe host fruit. At sympatric sites, the fruiting time of apple trees optimal for survival is earlier than hawthorns by ~3-4 weeks, and the life history timing of each host race is divergently adapted to these temporally distinct resources (Dambroski et al. 2007). The corresponding difference in the eclosion timing of the short-lived adult flies drives strong, but incomplete temporal reproductive isolation between the host races (Feder et al. 1994). The combination of temporal isolation and prezygotic reproductive isolation due to divergent chemosensory adaptation (Linn et al. 2003; Dambroski et al. 2005) limits on-going hybridization between apple and hawthorn flies at sympatric sites to 4-6% (Feder et al. 1994). Genetic differentiation in this system is based on consistent frequency differences in shared alleles rather than fixed variants between host races (Michel et al. 2010, Powell et al. 2013, 2022, Meyers et al. 2020).

Both populations of flies are attacked by three species of specialist parasitoid wasps in the family Braconidae: *Diachasma alloeum*, *Diachasmimorpha mellea*, and *Utetes canaliculatis* (Hood et al. 2015), abbreviated Da, Dm, and Uc, hereafter. Collectively, the flies and wasps represent one of the best supported cases of “sequential’’ ecological speciation, where the initiation of speciation via divergent selection at one trophic level triggers the subsequent initiation of speciation at higher trophic levels (Forbes et al. 2009; Hood et al. 2015). Moreover, each wasp species is also temporally specialized within fly hosts via divergent life history timing, which generates reproductive isolation between wasps attacking different hosts (Hood et al. 2015). Seasonal timing of adult emergence in this system is largely governed by the regulation of diapause – an ecophysiological state of dormancy. The environmentally sensitive periods that affect the initiation, maintenance, and termination of diapause development mainly occur after the flies have formed their puparia (Ragland et al. 2009; Powell et al. 2020), and both flies and parasitoids spend >85% of their life cycles in diapause inside sessile, behaviorally inert puparia. Thus, we can manipulate seasonal temperature regimes in biologically relevant treatments across this insect community during this key life stage dictating phenology.

As univoltine temporal specialists, the ability of *Rhagoletis* and their parasitoids to initiate and maintain diapause in the face of developmentally permissive temperatures before winter is critical for adaptive life cycle timing (Kostal 2006, Hahn & Denlinger 2011). The initiation and initial pre-winter maintenance of diapause are physiologically facultative in these populations. However, responses to environmental cues are sufficiently variable such that some fly and wasp individuals express maladaptive phenotypes that either forgo diapause initiation completely or fail to maintain metabolic and developmental suppression in high temperature conditions (Fig. 1; Dambroski & Feder 2007, Ragland et al. 2009, Calvert et al. 2022). In nature, flies that express either phenotype are ecologically doomed; flies that forgo diapause eclose as adults when host fruits are no longer available, and flies with short diapause lengths ramp up their metabolism, exhaust nutrient stores, and die before overwintering.

**Figure 1.**
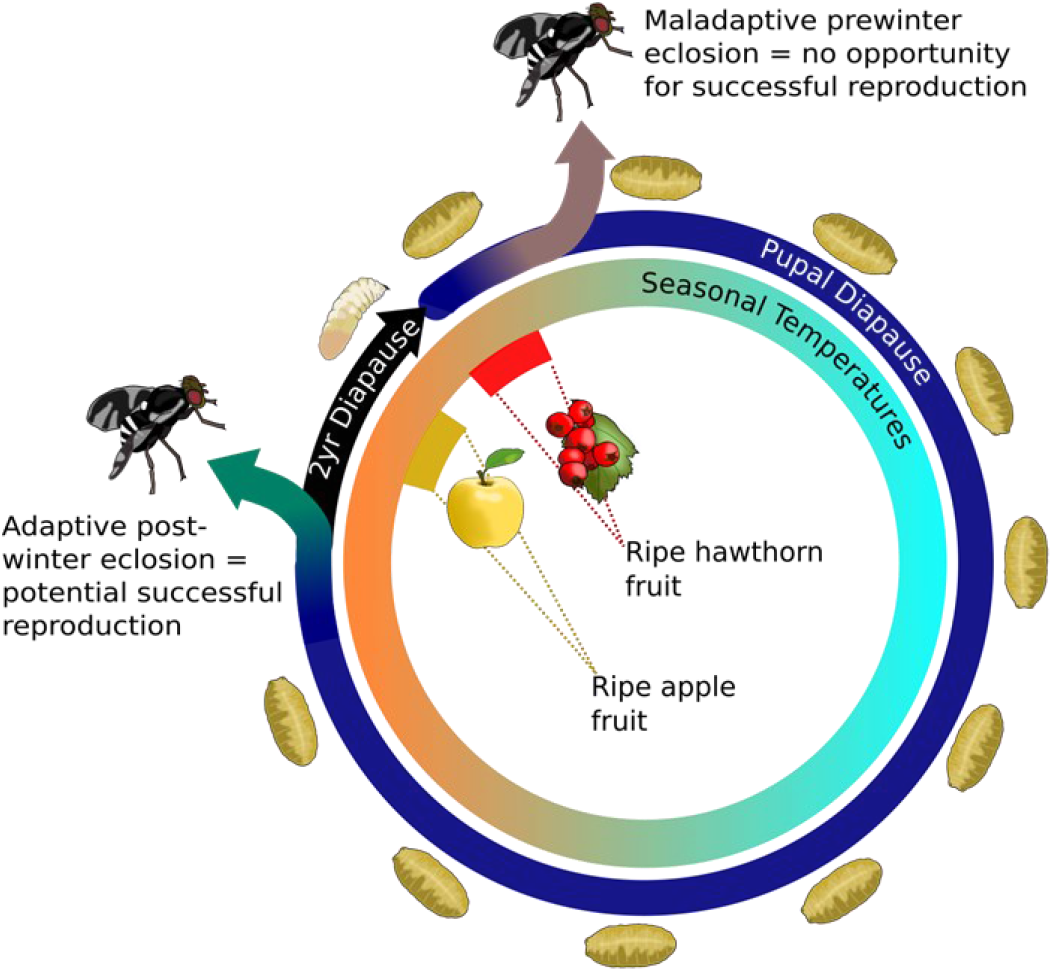
The lifecycle of *R. pomonella* flies, showing the potential for maladaptive pre-winter adult eclosion due to either a failure to enter pupal diapause or premature termination of diapause and the potential for multi-year diapause flies that remain in dormancy across the entire growing season.

In this study, we conducted a simulated climate change experiment on natural apple and hawthorn fly and parasitoid communities at a long-studied sympatric field site at Urbana, Illinois, USA (McPheron et al. 1988, Dambroski et al. 2005, Powell et al. 2020, Dowle et al. 2020). Insects were exposed to daily and seasonal temperature fluctuations reflecting either control (ten-year average of soil temperature) or elevated temperature (3°C warmer, see methods below). The tractability of this system allowed us to impose these experimental conditions across the entirety of the life stages affecting phenology in these species, including the initiation and termination of diapause. We tested how realistic simulated warming affected both flies and wasps in the community at three different levels: I) fitness changes based on maladaptive pre-winter diapause initiation and maintenance and phenological shifts within populations, II) phenological synchrony of communities across trophic levels, and III) the strength of prezygotic reproductive (temporal) isolation between divergently adapted populations.

## METHODS

### Field collections and insect husbandry

In August and September of 2017, we collected apple and hawthorn populations of *R. pomonella*, respectively, from infested fruit from wild populations in Urbana, IL during peak infestation for both plants. The apple and hawthorn sites are located ~1,300 m apart, well within the flight range of these flies (Roitberg et al., 1984). Field collection procedures followed long-established methods for *R. pomonella* husbandry (Feder et al. 1994, Dambroksi & Feder 2007, Powell et al. 2020) detailed in the supplementary methods. Infested fruit were collected from the ground and trees and transported to the laboratory where they were kept in an environmentally controlled room under standard *Rhagoletis* rearing conditions (25°C and 14:10 light: dark (L:D); Dambroski & Feder, 2007). Fruit were placed in wire mesh baskets in plastic trays. Wandering third-instar larvae emerging from the fruit dropped into the trays and were collected daily. Each day’s collection of larvae was equally divided and randomly assigned to control or warmed treatments. Larvae were subsequently maintained in Percival (Perry, IA) I-41-VL environmental chambers operating with C9 Intellus control systems, where they underwent pupal metamorphosis and experienced conditions simulating current versus predicted future temperatures, respectively (see below). Pupae were placed in individual tubes to track developmental timing from pupation to eclosion. A total of 1885 and 1719 apple puparia and 1453 and 1422 hawthorn puparia were monitored daily for adult fly and wasp eclosion in the control and warming treatments, respectively. Eclosing insects were sexed and wasps identified to species. At the end of one year, unenclosed puparia were frozen at −20 °C before being dissected under a stereomicroscope to determine the identity and developmental state of each insect.

Seasonal and daily temperature fluctuations for the control treatment were based on 10-year averages from 2007-2016 of 10 cm soil temperature data from the NOAA weather station at Watseka, IL (40.79, −87.76) closest to the Urbana, IL site (Fig. 2). Seasonal temperature changes were imposed over 1-week increments, and daily temperature fluctuations followed a periodicity based on the local seasonal photoperiod cycle. The circadian cycle for each day contains four set-points: the midpoint temperature at sunrise and sunset, the maximum temperature halfway between sunrise and sunset, and the minimum temperature halfway between sunset and sunrise, with the program set to ramp temperature linearly through the intervals between set points. Daily fluctuating conditions were maintained until average daily soil temperatures dropped below 6 °C, the physiological limit of *R. pomonella* development (Neilson 1962), at which point chamber temperatures ramped between a midpoint of 3.0 °C, maximum of 3.5 °C, and minimum of 2.5 °C until soil temperatures reached 6 °C again in the Spring. The simulated climate warming treatment was established as a 3 °C increase over the weekly temperature regimes of the control treatment (Fig. 2) based on projections for the Midwest for 2050 to 2100 for high to moderate IPCC emissions scenarios, respectively (IPCC 2013, Kunkel et al. 2013, Pryor et al. 2013). Thus, the warming treatment represents an aggressive model of climate change in the near term (<50 years) but a conservative model by the end of the century. Both temperature programs used sunrise and sunset times at the Watseka, IL station in 2016 to calculate daylengths.

**Figure 2.**
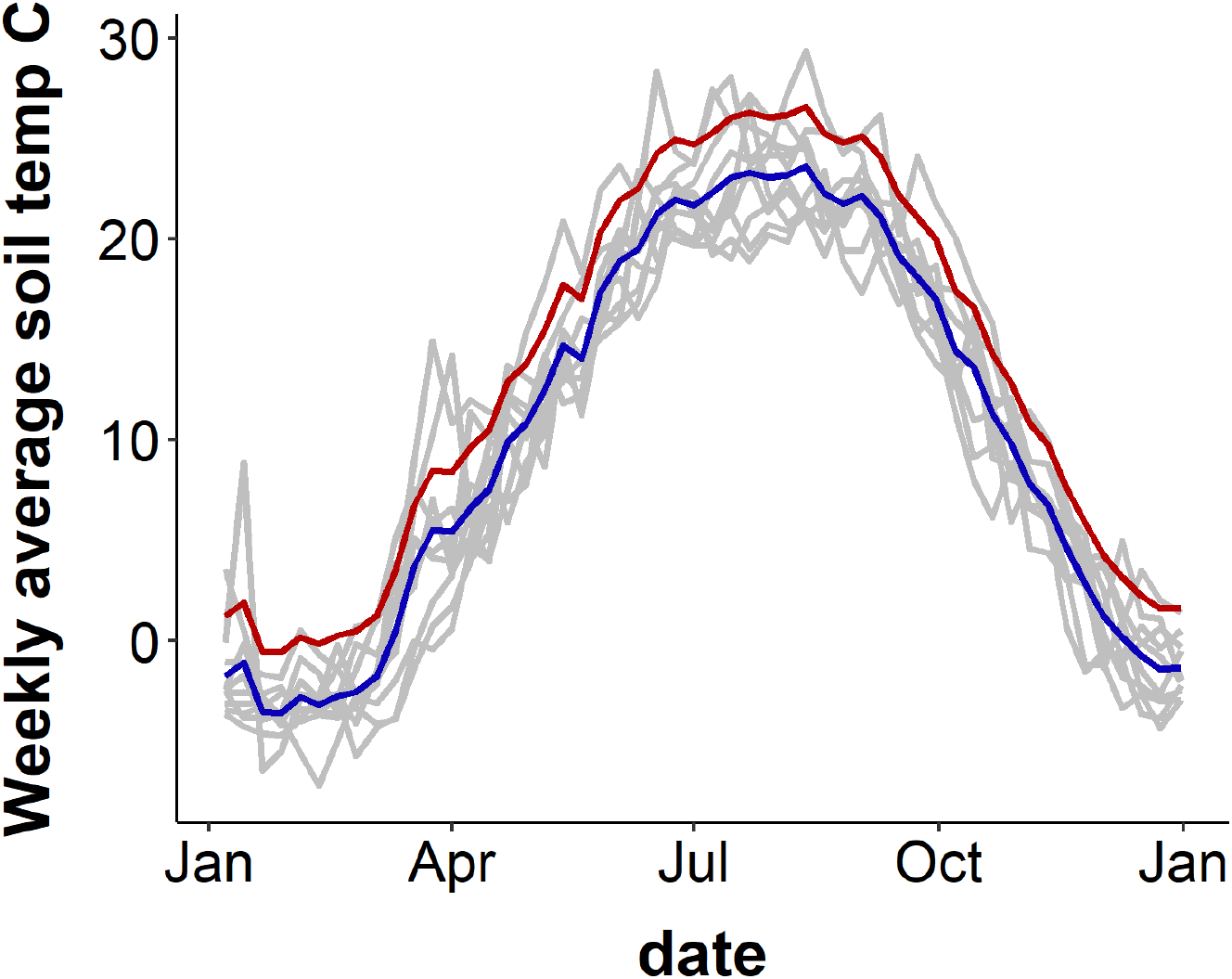
Weekly soil temperature averages in Celsius across the year. Control temperatures are shown in the blue line, and Warming temperatures are shown in red. Grey lines show each year from 2007-2016.

### Statistical analysis

Statistical analyses were performed in RStudio version 1.1.463 (RStudio Team 2022) using packages *survival* (Therneau & Grambsch 2000), *survminer* (Kassambara & Kosinski 2018), *ggplot2* (Wickham 2016), and *MASS* (Venables & Ripley 2002). To test for differences in phenology, we used accelerated failure time analysis, which is a type of survival analysis that uses survival regression models to test for differences in the timing of the probability of events (Fox 2001). We tested which variables explained variation in the timing of eclosion, and we describe model terms below. Models were fit with a loglogistic distribution, which was a significantly better fit than other distributions (loglikelihood tests for difference in fit compared to Weibull, Gaussian, logistic, lognormal: all X^2^ > 1064, all p < 0.0001).

First, we tested for differences in eclosion timing between control and warming temperatures within populations. For flies, the initial model included fixed effects for host plant, temperature regime, sex, and all interactions. We used AIC reduction in MASS to reduce initial models. The final model included main effects of host plant, temperature regime, and sex, as well as interaction terms between host plant and temperature treatment, and between host plant and sex. For wasps, the initial model was the best fit model, and it included fixed effects for host plant, temperature treatment, sex, and all interactions. To determine fitness changes between control and warming treatments, we tested for differences in the proportions of outcomes within flies and wasps using Chi-square tests. In flies, possible outcomes were: eclosed pre-winter, eclosed post-winter, remained alive as pupa (multi-year diapausing flies), died as pupa, and died as pharate adult prior to eclosion. In wasps, possible outcomes were: eclosed pre-winter, eclosed post-winter, died as larva, died as pupa, and died as pharate adult.

Second, we tested for differences in eclosion curves between flies and wasps reared in the same temperature treatment. The initial model was the best fit model, and it included fixed effects for insect type (fly or wasp), host plant, temperature treatment, and all interactions. We fit a separate model because we added a fixed effect for insect type to test for differences in response to temperature between flies and wasps. We did not include sex as a fixed factor because we did not hypothesize interactions between insect type and sex and because sample sizes of wasps of each sex were not large enough to test interactions between sex and other fixed effects. Third, we calculated the magnitude of temporal premating reproductive isolation (RI) between apple and hawthorn flies in the respective temperature treatments, accounting for effects of sex on eclosion time using a model of “co-occurrence” premating isolation from Sobel & Chen (2014). We used the number of flies from each population that eclosed on the same day (assuming all eclosing flies survived to reproductive maturity). We accounted for differences in relative population sizes using Sobel and Chen’s equation RI_4S2_ (2014). We also extended this equation to account for potential differences in eclosion timing of each sex by calculating RI from the perspective of each sex from each population (Fig. S1). We then averaged RI for each sex and each population within each temperature treatment to calculated total temporal isolation under current versus predicted climate change temperatures.

## RESULTS

### Phenological responses of flies

The apple and hawthorn fly populations differed in their pre-winter diapause initiation and maintenance responses to warming. No hawthorn fly eclosed before simulated winter in the control or warming temperature treatment (Fig. 3). In contrast, 4.6% of apple flies eclosed pre-winter in the control treatment, while 24% eclosed in the warming treatment, which is over a 5-fold increase (X^2^ = 281, df = 1, p < 0.0001, Fig. 3). Importantly, the prewinter eclosion times of all apple flies were later than the peak mating and oviposition time of the sympatric hawthorn population at Urbana. The earliest prewinter apple fly eclosed on September 16, while peak mating and oviposition in hawthorn flies occurs in late August to early September. *Rhagoletis pomonella* adults take 10-14 days after eclosion to sexually mature (Prokopy & Bush 1973), pushing predicted reproduction of prewinter eclosing apple flies into early October. Thus, even if some of the pre-winter apple adults were receptive to using hawthorn as a host (Forbes et al. 2005), the temporal window of availability for hawthorns would have already closed.

**Figure 3.**
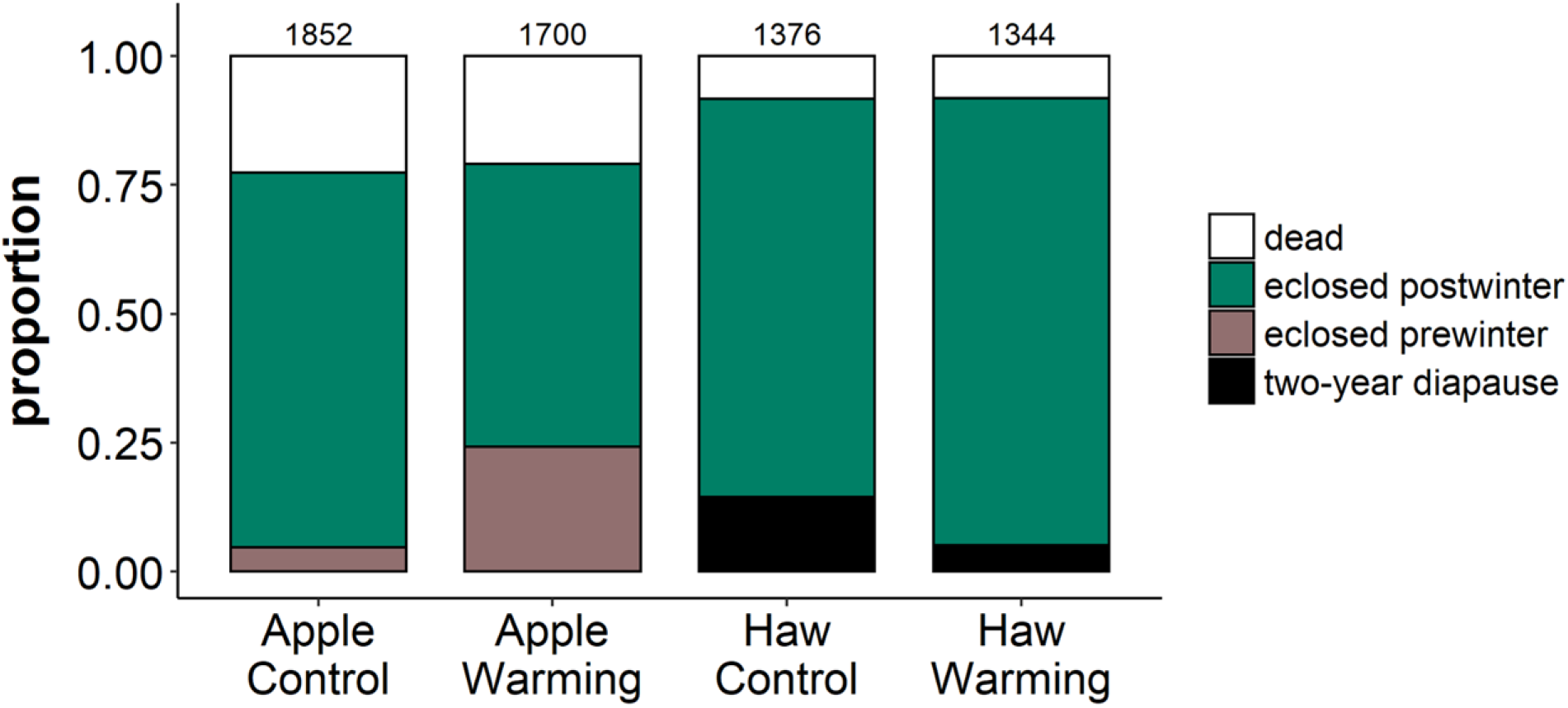
The fates of *R. pomonella* pupae from the apple and hawthorn (Haw) host races for both control and warming conditions with total sample sizes at the top of the bars. Proportions exclude parasitoid wasps, identified by successful adult eclosion or in post-mortem dissections.

While simulated warming did not induce non-diapause development in hawthorn flies, it did affect another aspect of diapause. Typically, a subset of *R. pomonella* flies eclose after two or more yearly cycles of chilling and heating rather than one (Phipps & Dirks 1933; Dambroski & Feder 2007). Multi-year diapause generates a seed bank of pupae in dormancy, likely representing a bet hedge against annual variability in host fruit set (e.g., Menu et al. 2000, Moraiti et al. 2014). Under warming, significantly fewer hawthorn flies remained in diapause at the end of the experiment (control: 14.4%, warming: 4.9%, X^2^ = 66.37, df = 1, p < 0.0001, Fig. 3). In the apple flies, no multi-year class pupae were observed in the control or warming treatment (Fig. 3).

Taken together, the results suggest that the warming temperature treatment affected what could be considered the front end of the diapause phenotype in the seasonally-earlier apple race by inducing an increased portion of individuals out of their normal one-year life cycle into non-diapause development (Fig. 3). Alternately, the warming temperature affected the back end for the seasonally-later hawthorn race, by causing an increased portion of individuals to shift from a multi-to one-year cycle (Fig. 3). These shifts did not change overall survivorship in the experiment between control and warm treatments within apple or hawthorn flies (Fig 3). However, the reduced proportion of multi-year diapausing hawthorn flies and increased numbers of non-diapausing apple flies in the warm temperature treatment are both maladaptive in nature, having negative fitness consequences that may adversely affect the long-term persistence of local populations in the face of climate change.

Simulated warming also affected the timing of post-winter adult eclosion (Fig. 4). The best-fit AIC-reduced accelerated failure time model retained a significant main effect of temperature (z = 10.44; p < 0.0001), with eclosion curves shifting earlier for both fly populations (Fig. 4A). However, a significant population by temperature interaction term (z = 4.76, p < 0.0001) indicated that the shift was more pronounced for apple flies (Fig. 4A). In addition to shifting median eclosion earlier, the warm treatment also caused 59% and 85% increases in the variance of eclosion times in apple and hawthorn flies, respectively (Fligner-Killeen test for homogeneity of variances in non-normal data, both host associations: X^2^ > 69, df = 1, p < 2.2×10^-16^; Figs. 4B, C).

**Figure 4.**
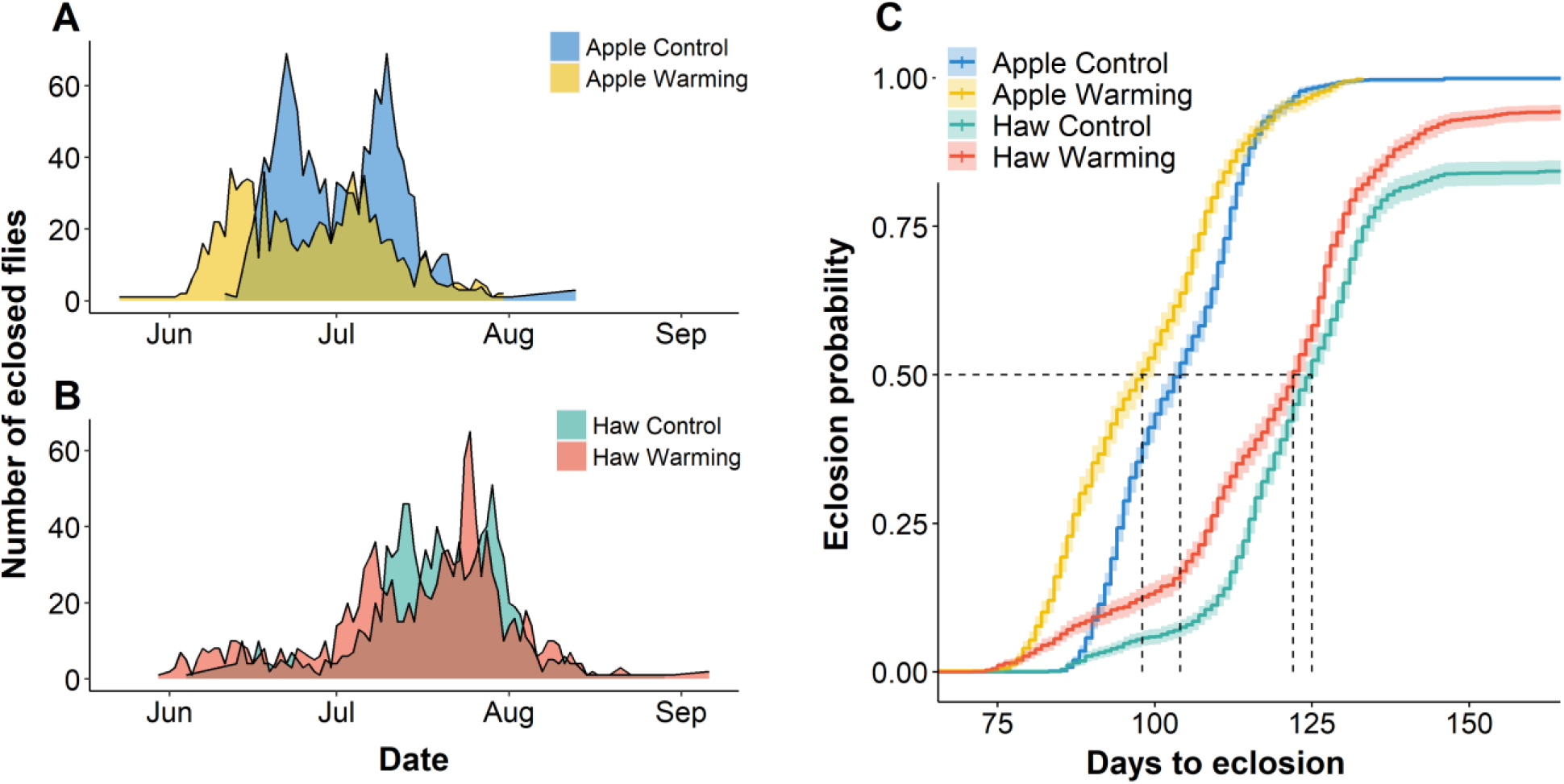
Post-winter adult eclosion distributions for (A) apple flies and (B) hawthorn flies. (C) Cumulative eclosion probability curves for apple and hawthorn flies reared under control and warming temperature regimes with days to eclosion since spring equinox, with confidence envelopes representing 95 % CI from the best fit accelerated failure time model. Dashed lines show days to eclosion at median eclosion probability for each population and temperature regime.

The asymmetric shifts in post-winter eclosion time of the apple and hawthorn flieshave implications for on-going divergence in this incipient speciation system. The change in post-winter eclosion distributions under warming conditions led to greater overlap of sexually active adults (i.e., increased potential for gene flow) in the warming compared to control treatment (Fig 6A, B). We calculated temporal RI to be 0.44 (95% CI: 0.31-0.56) in control and 0.33 (95% CI: 0.23-0.43) in the warming treatment, a 25% reduction.

**Figure 5.**
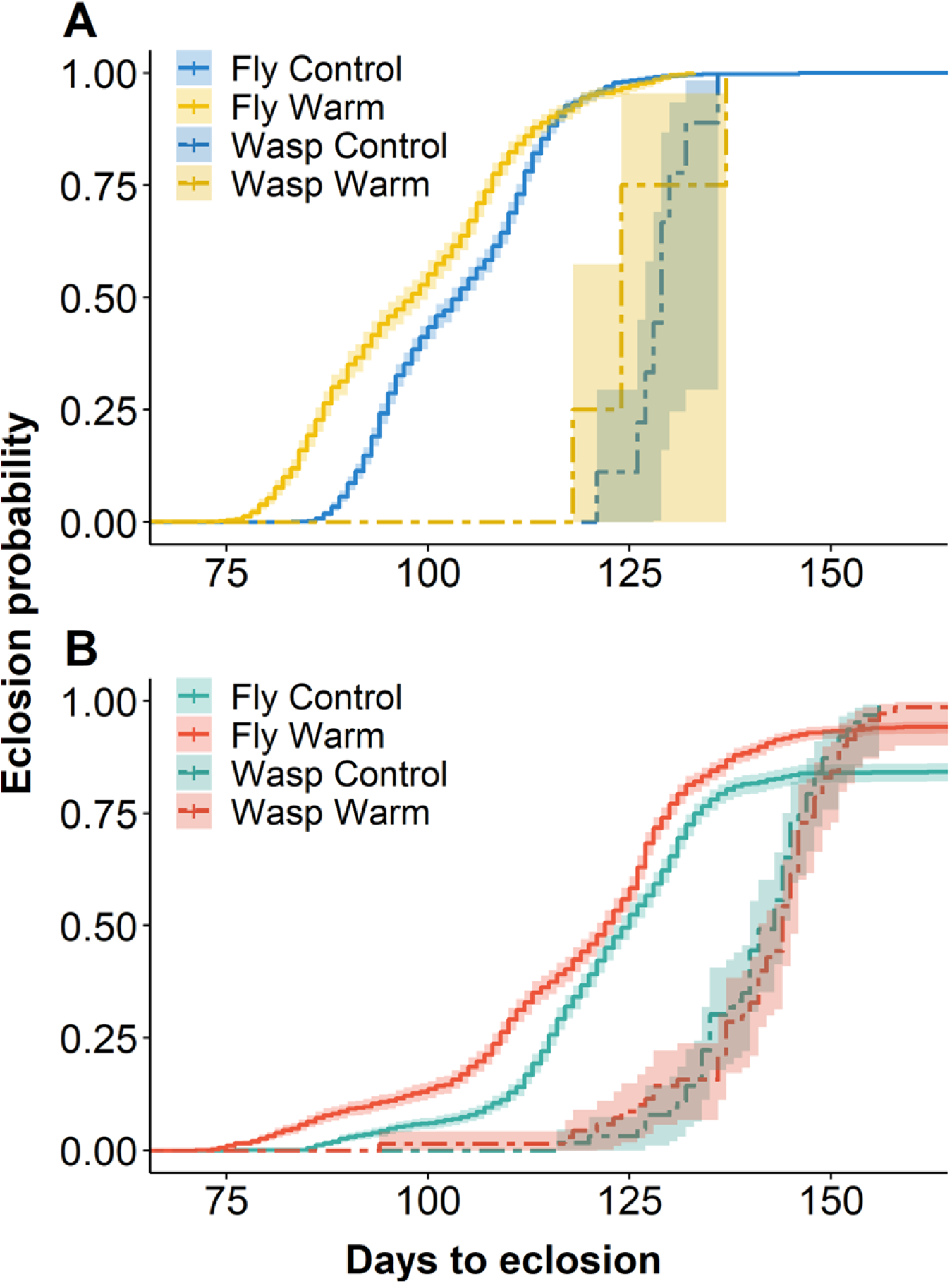
Cumulative eclosion probability curves for both flies and parasitoids reared from (A) apple and (B)hawthorn control and warming temperature regimes with days to eclosion since spring equinox, with confidence envelopes representing 95% CI from the best fit accelerated failure time model.

**Figure 6.**
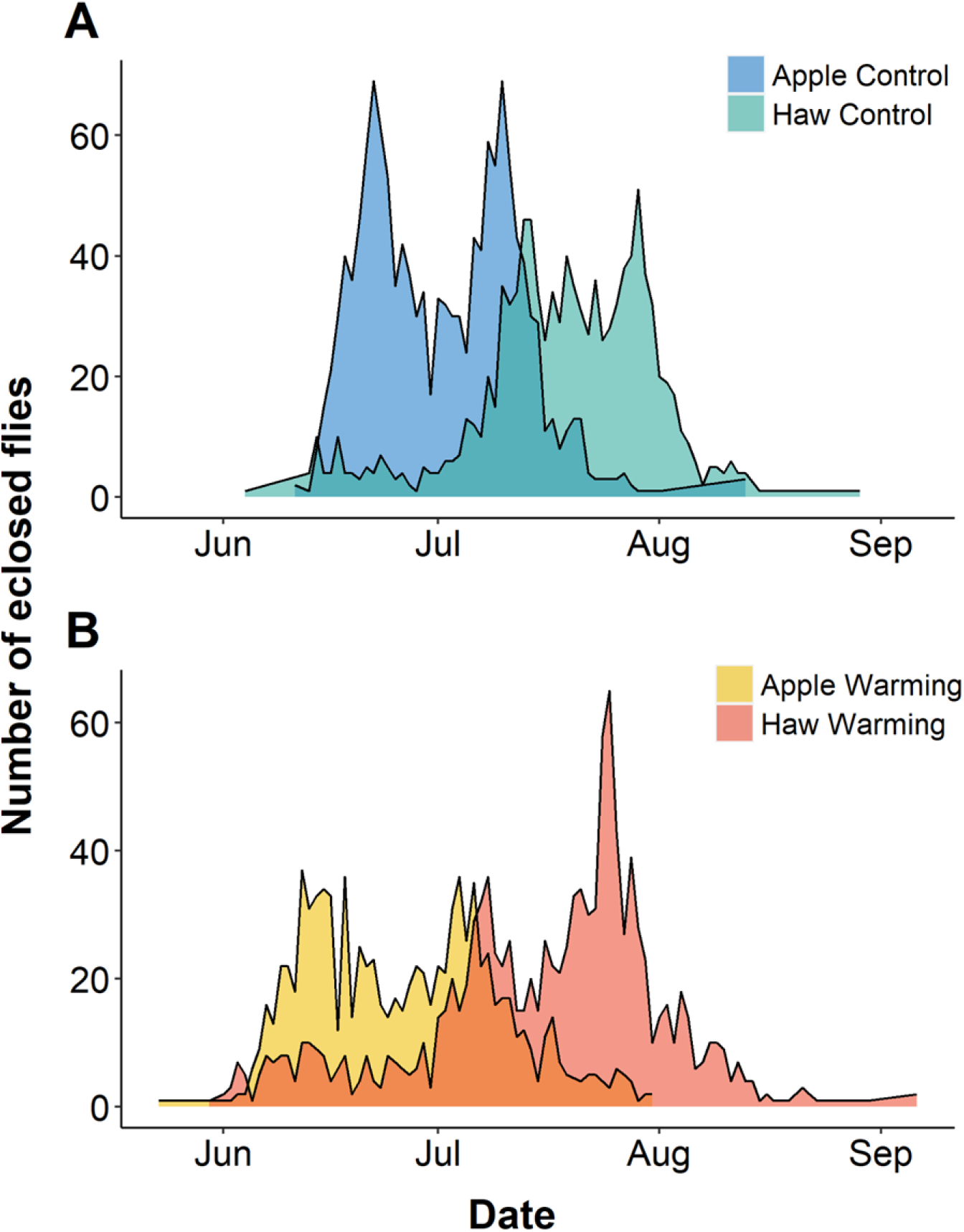
Post-winter eclosion distributions for apple and hawthorn flies in (A) control and (B) warming conditions, showing the overlap in phenology that determines the magnitude of temporal pre-zygotic reproductive isolation between the two host races.

### Phenological responses of parasitoids

Eclosion times of wasps did not differ between the control and warming treatments (Fig. 5), implying that parasitoids may not be as sensitive to increase temperature compared to flies. However, the sample size for apple fly parasitoids was small (n = 13), limiting our ability to detect significant differences if any exist. For hawthorn wasps (n = 134), accelerated failure time eclosion probability curves for control and warming treatments crossed each other and had overlapping 95% confidence intervals across the range of eclosion dates (Fig. 5). Thus, the curves showed no indication of separation at any point across their distributions (Figs. 5B). We also examined accelerated failure time eclosion probability curves for the three hawthorn wasps separately and they overlapped for each taxon between control and warming treatments (all model effect of temperature *β* < |0.02|, all p > 0.2, Figs. S2A-C). Thus, the lack of eclosion time response was not specific to a particular wasp but was general for all parasitoids. To examine our statistical power to detect a shift in eclosion due to warming in hawthorn wasps, we performed a permutation analysis in which eclosion times for hawthorn wasps were simulated to be shifted a day or more earlier in the warming treatment than the control (see online supplementary information). Our observed experimental results showed a weaker response coefficient than 94.4% of coefficients simulated from a 1-day shift and 98.9% from a simulated 2-day shift (Fig. S3). Thus, if the warming treatment induced a modest shift in eclosion time in the wasps, obscured by sampling error in our experiment, our observed results are still much weaker than would be expected by chance.

Combining eclosion data across temperature treatments showed that the three hawthorn wasp species differed in their eclosion phenologies (F = 18.75, p = 7.7 × 10^-8^, Fig. S4), as expected from previous research demonstrating temporal niche partitioning within the community (Hood et al. 2015, 2021). The relative order of eclosion of the wasps from earliest to latest was Uc, Dm, and Da. Moreover, like their fly hosts, all three species of parasitoid wasps are physiologically capable of forgoing diapause and entering direct adult development under prolonged periods of pre-winter heating in the laboratory (Hood 2015). However, no apple or hawthorn parasitoid in the control or warm treatment eclosed before winter.

## DISCUSSION

### Fitness consequences within fly populations are asymmetric

Despite diverging only in recent historical times (Bush 1966) and experiencing the same abiotic factors in sympatry, the apple and hawthorn host races, as well as their parasitoids, showed asymmetric responses to simulated climate change. These differences affecting propensities for non- and multi-year diapause development, and eclosion time, highlight how difficult it may be to generalize responses to climate change across not only species but conspecific populations. The regulation of diapause is driven by complex gene-by-environment interactions, involving the integration of multiple environmental cues with multiple physiological systems and gene co-regulatory networks (Emerson et al. 2009; Dowle et al. 2020). Thus, how climate change affects this critical feature of seasonal adaptation in insects may depend on population-specific thermal reaction norms underlying the regulation of different phases of diapause development. While the regulation of diapause development is the major determinant of adult eclosion phenology in *Rhagoletis* (Powell et al. 2020), assessing how other life stages, e.g., larval development within fruit, sexual maturation, and longevity may be affected by warming will give a more complete picture of the fitness consequences of altered seasonal regimes.

### Interactions across trophic levels

The phenological shift in the flies coupled with the apparent lack of shift in the parasitoids suggests that climate change may cause phenological asynchrony across trophic levels. All three parasitoid species have shorter adult lifespans than the flies (<10 days vs. ~30 days; Forbes et al. 2009; Hood et al. 2015). Thus, an earlier shift in fly phenology of just a few days without a compensatory shift in the wasps could result in substantially decreased availability of hosts at the right life stages for successful parasitoid attack (Hood et al. 2015). Reduced top-down control could allow fly populations to increase, potentially intensifying competition within populations for host plant resources. However, shifted phenology in fly populations will also generate stronger selection on wasps to match a new phenological optimum. The strength and direction of such selection depends on whether host-parasitoid phenological relationships are currently optimal from the perspective of the parasitoids (Singer & Parmesan 2010). Gene x environment interactions affecting variation in eclosion in the parasitoids have been less well studied than those of the flies (Dambroski & Feder 2007). However, eclosion time in the wasps has been shown to be genetically based and polygenic (Forbes et al. 2009; Hood et al. 2015), like the flies (Feder et al. 1993; Doellman et al. 2019), and, thus, parasitoids may have the capacity for rapid adaptation, tracking phenological shifts in their host flies.

Simulated warming had different effects on both the position and shape of adult eclosion phenology distributions between apple and hawthorn flies (Fig. 4). How these differences manifest as fitness consequences and how natural selection may act on those fitness regimes under future conditions will also depend on how the fruiting phenology of its hosts are affected by the same conditions. While an experimental test of simulated climate change on the apple and downy hawthorn trees in Urbana, IL is logistically unfeasible, previous empirical and theoretical work on apple and other fruit trees in Rosacea speak to possible effects of predicted future conditions on host fruit phenology. Studies of apple phenology over the past several decades have shown a pattern of earlier flowering and fruiting through time (e.g., Wolfe et al. 2005; Fujisawa & Kobayshi 2007). However, models of apple phenology predict that long-term trends heading into an even warmer climate may be more complicated, due in part to antagonistic effects of warming during different seasons (Darbyshire et al. 2017). While warmer springs accelerate apple flowering times, shorter, milder winters have an opposing effect on phenology; thus, models predict advancing, stationary, or even receding timing depending on localized climate projections (Darbyshire et al. 2014). Moreover, such models predict that the effects of future climatic conditions on apples will be highly variable across geography and apple cultivars (Legave et al. 2015). The drivers of phenological variation in downy hawthorn have not been studied directly. However, more distantly related Rosacea fruit trees, such as *Prunus sp*., appear to show similar patterns of high intraspecific variation in phenological responses to seasonal temperature regimes, with contrasting effects of winter length and spring temperatures (Doi et al. 2010, Castède et al. 2014). In the midwestern United States, where our study site is located, latitudinal variation in fruiting phenology for apple and downy hawthorn trees runs in opposite directions, with apples fruiting earlier and hawthorns later moving North to South (Feder et al. 1989, Dambroski & Feder 2007). Thus, as the prevailing seasonal regimes become warmer and more erratic in coming decades, the differences in post-winter phenology effects between apple and hawthorn flies observed in our experiment may be superimposed on highly variable effects on fruiting phenology, both between their respective host plants and across geographic sites.

### Phenological shifts reduce temporal isolation between fly populations

Shifts in phenology due to warmer rearing temperatures lead to predicted greater temporal overlap of apple and hawthorn flies, with a 25% reduction in the strength of temporal isolation. Because temporal isolation is an early acting form for premating isolation, it can often have disproportionate effects on total RI compared to later acting barriers (Ramsey, Bradshaw, & Schemske 2003). Cases of incomplete ecological speciation may exist in a fragile equilibrium between migration and divergent selection (Nosil et al. 2009). Thus, increased migration stemming from weakened temporal isolation between apple and hawthorn flies could lead to increased genetic homogenization and even the possible collapse of differentiation between the host races.

However, increased seasonal overlap between the flies is ultimately dependent on convergence in the fruiting phenologies of apple and hawthorn trees. Further study is required, but as discussed above, a balance between the effects of warmer growing seasons and winters may result in host fruiting times remaining similar or even diverging under future conditions. If true and given the extensive standing genetic variation for life history traits in *R. pomonella* (Michel et al. 2010; Doellman et al. 2018), then divergent selection on eclosion timing in F1 offspring could counter increased migration rates in the parental adult generation, potentially maintaining or strengthening current levels of temporal RI. Our results highlight the precarious environmental sensitivity of temporal isolation. The increased frequency of extreme heat anomalies expected under climate change (Cai et al. 2014) may lead to more frequent and stronger oscillations in genetic divergence between apple and hawthorn flies and their associated parasitoids in coming years.

Certain populations of *R. pomonella* may be well poised for rapid adaptation to climate change. Standing variation in the form of latitudinal clines in diapause phenotypes (Dambroski & Feder 2007) and chromosomal inversions associated with these traits exist in both host races (Feder et al. 2003; Michel et al. 2010; Doellman et al. 2019). These clines allow apple and hawthorn flies to track shifts in host phenology. Just as poleward range shifts of whole species may be an important response to a warming world (Pelini et al. 2009), shifting positions and slopes of genetic clines for dormancy traits may be another important consequence for systems with geographic variation in dormancy traits (Bradshaw & Holzapfel 2006; 2008). As the genetic differences between the host races are superimposed over these clines (Feder et al. 1990), these shifts in the shapes of diapause responses to temperature may have further consequences for patterns of divergent selection and gene flow between apple and hawthorn flies in the future. Notably, the apple population at Urbana, IL is near the southern range limit of apple flies (Bush 1969), potentially limiting the amount of standing variation available for adapting to even earlier phenologies and hotter conditions. Moreover, antagonistic genetic correlations among pre- and post-winter diapause traits associated with chromosomal inversions (Calvert et al. 2022) may pose serious constraints on adaptation to novel conditions in this system.

## Conclusions

The asymmetric responses in diapause development and timing we observed in the *R. pomonella* system imply that temperate insect communities may be particularly vulnerable to fitness declines, altered species interactions, and the breakdown of divergent local adaptation. Even species or populations that may appear physiologically inert to increased temperatures, such as the parasitoid wasps here, may suffer fitness consequences as the result of asynchrony with interacting species. The proximate effects of climate change may ripple across ecological communities in complex ways, involving not just demographics and the temporal synchrony of species interactions but potentially altering the evolutionary landscape of divergent adaptation and speciation. Ecological theory predicts that specialist taxa may be at greater risk to environmental instability than generalists (Futuyma & Merron 1988; Scheiner 2002). Importantly, response to climate change also predicts a potential loss of biodiversity in both parasitoid and fly populations, though via different mechanisms. In the parasitoids, potential phenological mismatch of hosts and the very narrow adult life stage of parasitoids threatens biodiversity through fitness declines and perhaps even local extinction in the parasitoids during hotter years. In the flies, the on-going evolutionary process of ecological speciation driven by divergent specialization may be at risk of collapse due to population-specific shifts in phenological distributions that erode temporal isolation. More broadly, our findings of complex sensitivity of life history timing traits to changing thermal regimes predict consequences not only for the maintenance of current biodiversity but also the potential for new species to evolve. The parameter space under which ecological speciation can proceed in sympatry is narrow (Nosil et al. 2009; Powell et al. 2014) and systems with environmentally sensitive isolating mechanisms may be pushed into unfavorable parameter space by climate change. Coupled with empirical evidence for long-term declines in insect abundance (Wagner 2020), the complex effects of simulated climate change demonstrated here portend that the coming decades may see multifaceted changes to on-going ecological and evolutionary processes in the most diverse group of organisms on Earth.

## Acknowledgements

We thank undergraduate researchers A. Ahmed, A. Dukat, W. Estevez, D. Fama, M. Molina Mera, N. Mirza, A. Murray, I. Pyatetsky, E. Romeo for invaluable help executing, maintaining, and monitoring this experiment, and G.R. Hood for helpful discussions and comments on the manuscript. The work was supported by National Science Foundation grants 1639005 to THQP and DAH, 1638951 to GJR, and 1638997 to JLF.

## SUPPLEMENTARY INFORMATION

### Permutation analysis of parasitoid phenology effects

We conducted a permutation analysis to determine whether our observed lack of significant phenology effects in hawthorn wasps (Fig. 5) was an artifact of the relatively lower sample size in parasitoids compared to flies. This approach asked how likely the lack of accelerated phenology we observed in the hawthorn wasps (main effect of temperature: β = 0.0096) was to occur if we simulated 1 – 8 day acceleration effect of the warming treatment. First, we produced permuted samples matching the sizes of our control and warming samples (n = 63 and n = 70, respectively) by sampling across all hawthorn wasps eclosion dates across treatments. For each simulated shift values (0 – 8), we subtracted that value from each eclosion date in the resampled warming (n = 70) group. For example, to simulate a 2-day accelerated phenology effect of warming, the distribution of the resampled wasps assigned to the “warming” group was shifted forward by 2 days. We then applied the same Accelerated Failure Time model of eclosion time as a function treatment to the simulated data to obtain the β estimate for the main effect of temperature treatment. We conducted 10,000 iterations of this for each simulated shift value to produce the distribution of β estimates, and then asked what proportion showed a stronger effect of warming on accelerated phenology. As the effect of accelerated phenology is to lower the days to adult eclosion, β estimates consistent with this direction of phenological shift are negative. Even without a simulated shift in the resampled values (shift of 0), 82.8% of the iterations showed lower coefficients of temperature effect than the results of our experiment (Fig. S3). For the shifts of 1 day, 94.4% of the simulations exceed the experimental results, followed by 98.8% for 2 days and 99.8% and 3 days (Fig. S2). For days 4 and above, 100% of the simulations exceeded the experimental results.

### Supplementary Figures

**Figure S1.**
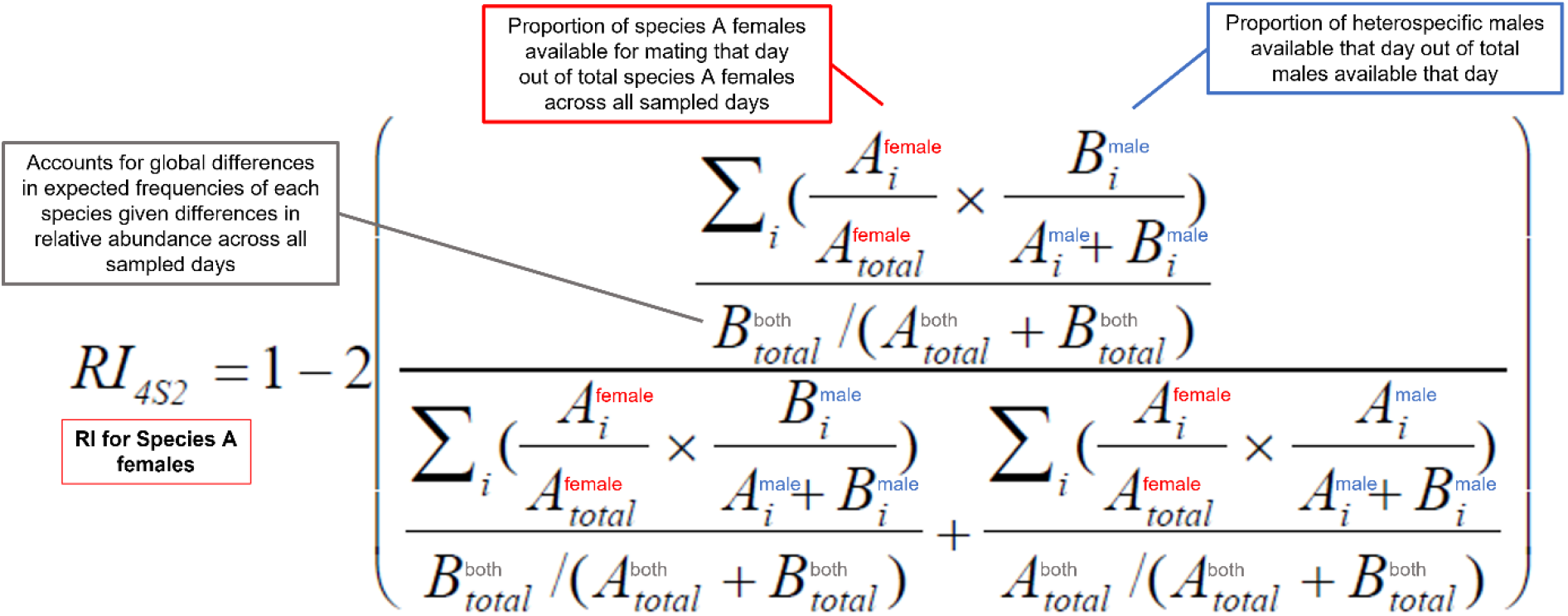
The RI_4S2_ formula from Sobel and Chen (2014) calculates RI for each species separately and accounts for differences in population sizes for each species. We revised this equation to calculate RI for each sex of each species separately. We label terms for females (red), males (blue), and both sexes combined (grey). In the numerator, we describe the components of the equation, and these components repeat with some differences in the denominator.

**Figure S2.**
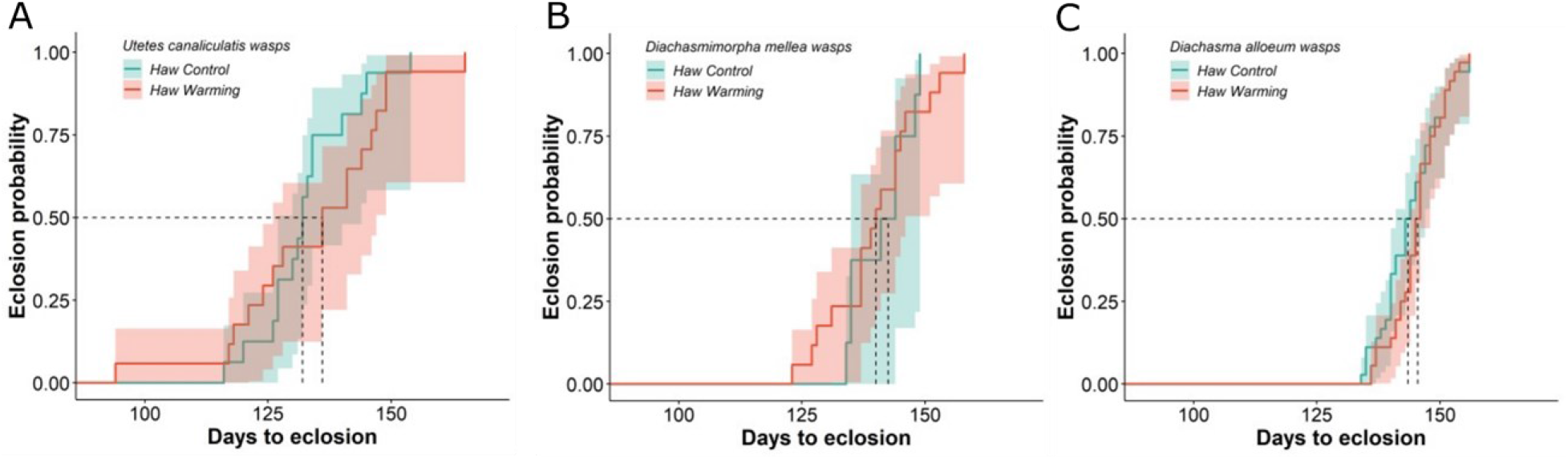
Cumulative eclosion probability curves for hawthorn-fly-infesting (A) *Utetes canaliculatis*, (B) *Diachasmimorpha mellea*, and (C) *Diachasma alloeum* parasitoid populations reared under control and warming temperature regimes as days to eclosion since spring equinox, with confidence envelopes representing 95% CI from best fit accelerated failure time models. Dashed lines show median days to eclosion for each species and temperature regime.

**Figure S3.**
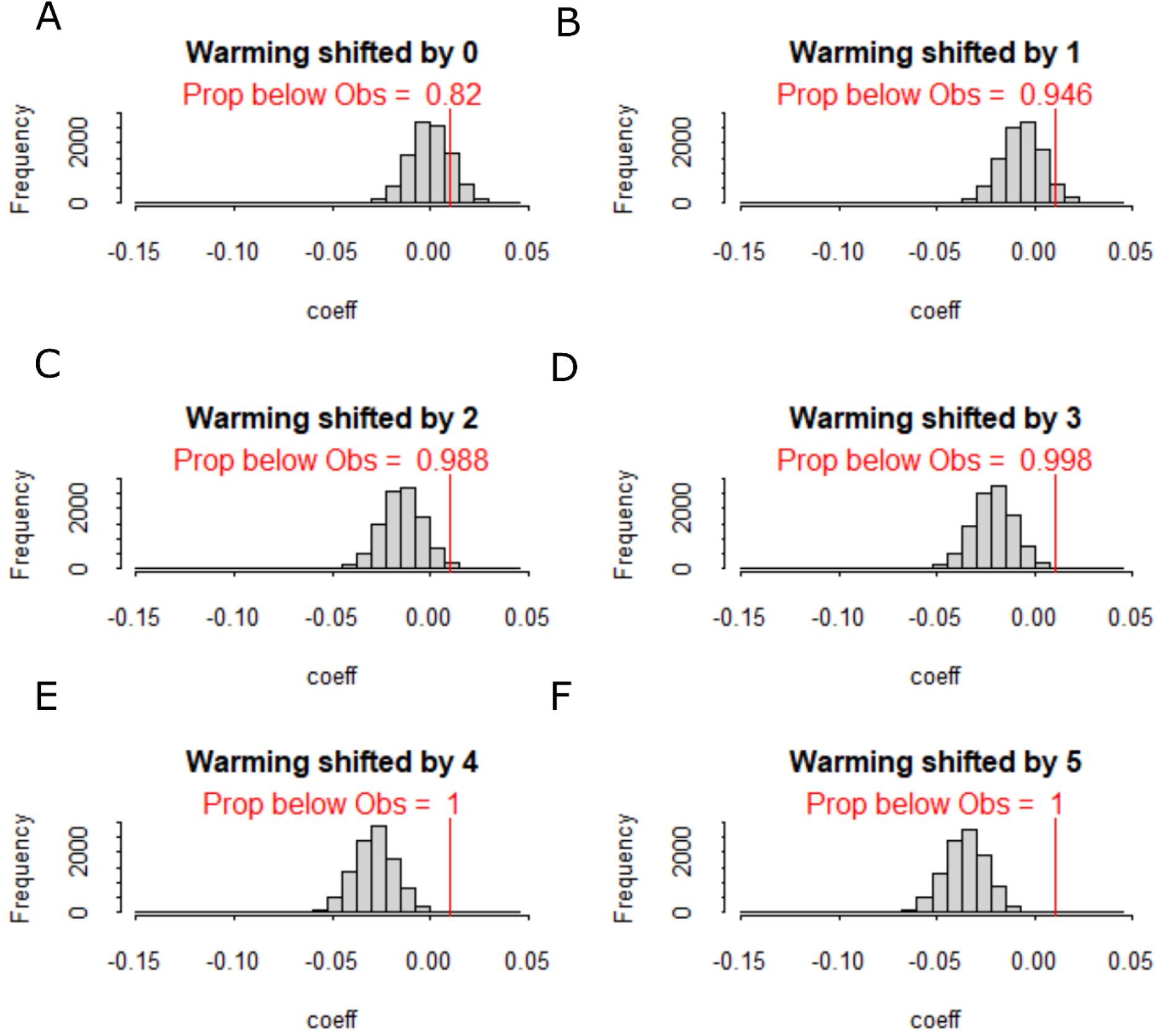
Distributions of accelerated failure time model coefficients of warming effects for 10,000 permutations of resampled parasitoid eclosion phenologies imposing (A) no shift and shifts of (B) one, (C) two, (D) three, (E) four, (F) and five days earlier in the simulated warming sample. Red line indicates the non-significant, slightly positive coefficient (β = 0.0096) in the observed data from the experiment. Red text describes the proportion of permutations that showed a lower coefficient than the observed value.

**Figure S4.**
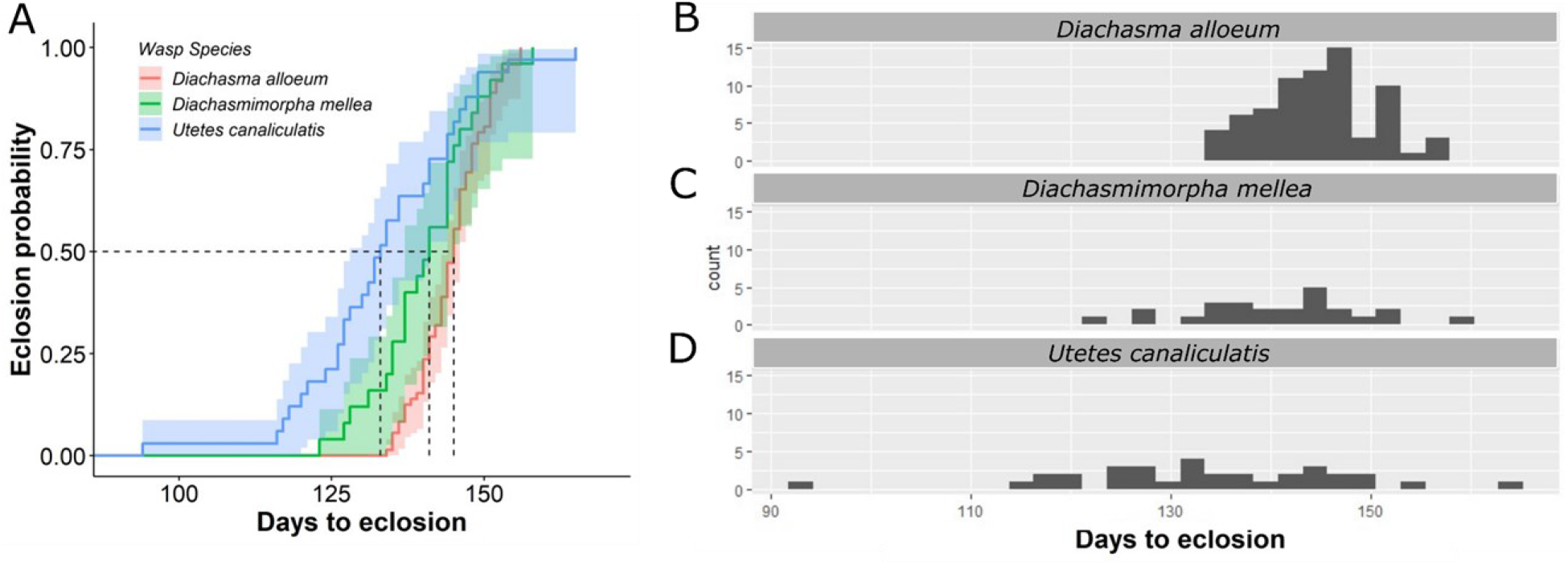
(A) Cumulative eclosion probability curves for *D. alloem* (red), *D. mellea* (green), and *U. canaliculatis* (blue) wasps across both temperature treatments with 95% confidence envelopes from accelerated failure time analysis, and distributions of observed eclosion timing for parasitoids across both treatments for (B) *D. allouem*, (C) *D. mellea*, and (D) *U. canaliculatis*, which was analyzed using a linear model framework to test for a main effect of species identity.

